# Shear Stress Induces Concentration Gradient Distributions of Membrane Proteins in Live Cells

**DOI:** 10.1101/2025.08.27.672548

**Authors:** Sawako Yamashiro, Misato Nomura, Nils Chapin, Sreeja Sasidharan, Louis Elverston, Leah Knepper, Damien Thévenin, Naoki Watanabe, Aurelia R. Honerkamp-Smith

## Abstract

Cells sense and respond to fluid shear stress. Cell surfaces are exposed to flow, yet the influence of shear stress on the behavior of plasma membrane proteins remains unclear. Here we show that extracellular flow induces the gradient distribution of cell membrane proteins with increasing concentration toward the downstream direction of the flow. Shear stress at 10-30 dynes/cm^2^ caused formation of concentration gradients of both GPI-anchored proteins and transmembrane proteins, including integrinß1, E-cadherin and the insulin receptor in *Xenopus* XTC cells. Using single-molecule live-cell imaging, we found that GPI-anchored T-cadherin molecules are dragged along the direction of flow under shear stress. In addition, shear stress induced concentration gradients of membrane proteins in COS-7 cells and human umbilical vein endothelial cells (HUVECs). Our findings suggest that external flow directly transports membrane proteins, establishing concentration gradients that may contribute to the cellular flow-sensing mechanism.

## Introduction

In living organisms, directional fluid flows occur both outside the cell (e.g., blood circulation, lymphatic flow) and within the cell (e.g., cytoplasmic streaming, the retrograde actin flow), serving various functions. Shear stress arising from blood flow constantly acts on the surface of the vascular endothelium, contributing to vascular homeostasis [1, 2] and cardiovascular development [3–5]. Disturbed blood flow in arterial vessels influence pathological processes such as in atherosclerosis [2, 6, 7]. Within the cell, the cytoskeleton-driven cytoplasmic flows are critical for distribution of organelles [8], mixing cytoplasmic contents [9], cell polarity formation [10], migration [11], and plant development [12]. To understand how flow functions, it is crucial to elucidate how flow forces influence the dynamics of molecules exposed to them.

Intracellular flow applies direct forces to molecules, causing them to move. The retrograde filamentous actin (F-actin) flow at the cell periphery causes downstream-biased gradient misdistribution of F-actin-binding probes including Lifeact and phalloidin [13]. The generation of molecular concentration gradient by intracellular flow has also been shown by mathematical simulations [13, 14]. Not only intracellular flow, but also extracellular flow may generate concentration gradients of molecules on the cell membranes. In model systems of glass-supported lipid bilayers, flow that generates low shear stress of 1-5 Pa (10-50 dynes/cm^2^) passively transports membrane-linked proteins over large distances to form micron-scale concentration gradients [15, 16]. These findings suggest that external flow can induce concentration gradients of membrane proteins along the flow direction within the upper plasma membrane of cells. Importantly, gradient distributions generated by lateral transport of membrane proteins may contribute to cellular responses to external fluid flow, such as determining cell polarity [17].

Several studies show that membrane proteins accumulate downstream of applied shear stress. The swimming of the unicellular parasite *Trypanosoma brucei* generates hydrodynamic forces that cause accumulation of the antibody-bound, GPI-anchored variant surface glycoprotein at the posterior of the cell [18]. Vascular endothelial-protein-tyrosine phosphatase (VE-PTP) accumulates at the downstream edge of cultured endothelial cells under shear stress [19] and in endothelial cells of the mouse aorta [20]. The study by Mantilidewi et al proposed that the polarized distribution of VE-PTP is caused by actin reorganization downstream of the shear stress-induced Cdc42 activation [19]. Shear stress also induces a downstream accumulation of heparan sulfate that covers the apical surface of cultured endothelial cells [21]. These studies support the hypothesis that exposure to fluid flow may induce the formation of a concentration gradient of membrane proteins on the surface of living cells. However, it is uncertain whether external flow directly transports membrane proteins across the plasma membrane in living cells. Furthermore, what types of membrane proteins may form the gradient distribution under shear stress remain unclear.

In this study, we show that extracellular flow-induced shear stress on cultured cells generates concentration gradient of membrane proteins along the flow direction. Under physiological conditions, shear stress in the human aorta is ∼10-20 dynes/cm^2^ ^[22]^. We show that shear stress at physiological levels induces movement of both GPI-anchored proteins and transmembrane proteins, including receptors and adhesion molecules. Single-molecule live-cell imaging under shear stress reveals that GPI-anchored molecule diffusion is biased in the direction of flow. These findings suggest that external flow directly transports membrane proteins, generating concentration gradients that may function in cellular mechanisms for sensing and transmitting mechanical information from external flow.

## Results

### Shear stress-induced gradient distribution of GPI-anchored proteins

We first noticed that fluid shear stress caused gradient distribution of GPI anchored EGFP (EGFP-GPI) in *Xenopus* XTC cells. The change in distribution of EGFP-GPI in cells exposed to shear stress was observed by fluorescence time-lapse microscopy. EGFP-GPI was almost uniformly distributed in XTC cells before application of shear flow (Fig. 1, A-D, Before). After applying shear stress to cells at 20 dynes/cm^2^ and 30 dynes/cm^2^, the fluorescence signal of EGFP-GPI was reduced at the upstream, and increased at the downstream (Fig. 1, A-D). In cells expressing EGFP without GPI, applying shear stress did not change the distribution of EGFP (Suppl. Fig. S1). When the external flow was stopped after the shear stress-induced EGFP-GPI gradient distribution was formed, the EGFP-GPI gradually returned to a uniform distribution (Suppl. Fig. S2). These results indicate that application of shear stress causes the formation of a concentration gradient distribution of EGFP-GPI that is anchored to the plasma membrane.

**Fig. 1.**
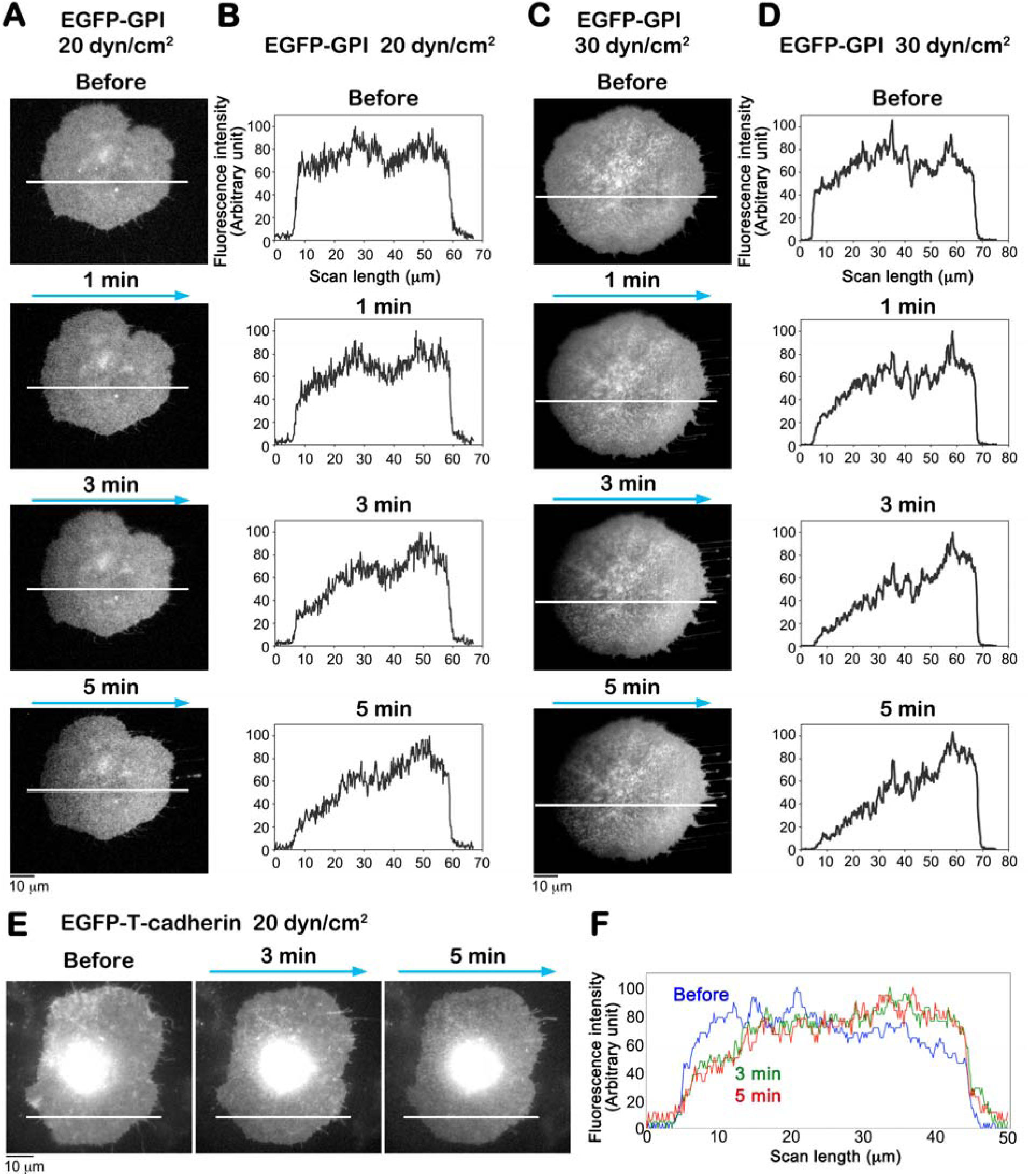
Extracellular flow induces the formation of a concentration gradient distribution of GPI-anchored proteins. A and C, The localization of EGFP-GPI expressed in XTC cells, before and after the application of shear stress in the direction indicated by arrows. B and D, Linescan analysis performed along the white lines in A and C. E and F, The formation of EGFP-T-cadherin concentration gradient under shear stress (20 dyn/cm^2^). A-F, The experiments were repeated at least 3 times and representative data are shown. Bars = 10 μm

We compared the fluorescence intensity of EGFP-GPI between the upstream and downstream sides of the cells. The ratio of the average value within 10 µm from the downstream cell edge to the average value within 10 µm from the upstream cell edge was 0.960 for before, and, 1.27, 1.99 and 2.49 for 1, 3, and 5 min after application of shear stress at 20 dynes/cm^2^, respectively (Fig. 1, B). Similarly, the ratio of the average fluorescence intensity at the downstream region to those at the upstream region was 1.19 for before, and, 1.93, 4.08 and 5.71 for 1, 3, and 5 min after application of shear stress at 30 dynes/cm^2^ (Fig. 1, D). These results indicate the following two points: (1) Shear stress induces gradient distribution of EGFP-GPI in the order of minutes. (2) The higher the shear stress magnitude, the steeper and faster the concentration gradient of EGFP-GPI is formed.

Next, we examined whether EGFP-fused T-cadherin forms a gradient distribution upon shear stress. T-cadherin is a GPI-anchored non-classical cadherin [23], and is highly expressed in blood vascular endothelial cells [24]. Similar to EGFP-GPI, shear stress at 10-30 dynes/cm^2^ induced a concentration gradient distribution of T-cadherin in the order of minutes (Fig. 1, E and F, Suppl. Fig. S3). These results suggest that extracellular flow at physiological magnitude, 10-20 dynes/cm^2^ [22], could cause the formation of a gradient distribution of GPI anchored membrane proteins.

### Fluorescent single-molecule analysis of T-cadherin under shear stress

To elucidate how extracellular flow affects the behavior of GPI anchored proteins, we performed fluorescent single-molecule imaging of T-cadherin under shear stress. We used the T-cadherin probe, in which SNAP-tag was inserted into the 4^th^ and 5^th^ extracellular cadherin repeats (EC domains) of T-cadherin (Fig. 2, A). XTC cells expressing the T-cadherin probe were treated with the cell-impermeable fluorescent substrate SNAP-surface549 (NEW ENGLAND BioLabs) to label the cell membrane-bound T-cadherin probe. The SNAP-surface549 labeled probe was suitable for fluorescent single-molecule microscopy in terms of brightness and photostability. We acquired continuous images with an exposure time of 50-200 ms, without intervals, for 20-60 s before and after applying shear stress (Fig. 2, B).

**Fig. 2.**
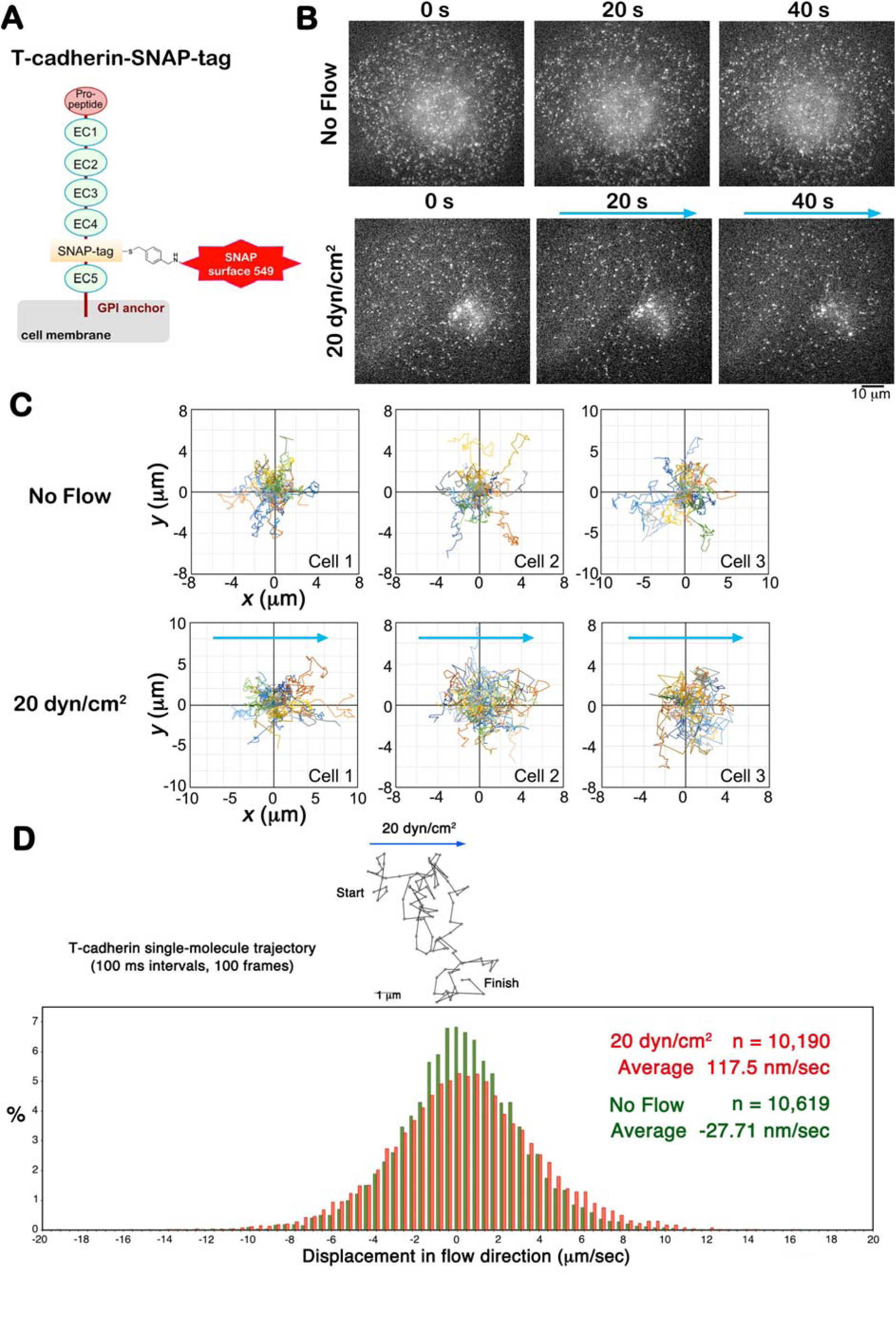
Fluorescent single-molecule analysis of T-cadherin under shear stress. A, The SNAP-surface549 labeled T-cadherin probe for fluorescent single-molecule imaging. B, Fluorescence single-molecule images of SNAP-tag T-cadherin in XTC cells with or without extracellular flow. Series of fluorescent T-cadherin single-molecule images were acquired with a 200 ms exposure time without intervals. C, Trajectories of T-cadherin molecules (10 sec, 50 frames) in each of the three cells under control (Cell 1, n = 25; Cell 2, n = 25; Cell 3, n = 22) or shear stress at 20 dyn/cm^2^ (Cell 1, n = 24; Cell 2, n = 30; Cell 3, n = 22). The starting point of each trajectory is plotted at the intersection of the X and Y axes. The arrow indicates the flow direction. D, Displacement distributions of T-cadherin single-molecules in no-flow control cells (green) and cells under shear stress at 20 dyn/cm^2^ (red). The average value of the displacement is increased in cells under shear stress.

First, we compared trajectories of T-cadherin single-molecules for 10 s between under the control condition without external flow and under shear stress at 20 dynes/cm^2^. For the condition under shear stress, T-cadherin single-molecules were tracked within 1 to 2 mins after applying external flow. In the control XTC cells, T-cadherin single-molecules moved without bias in the directionality of its diffusion (Fig. 2, C, upper), whereas T-cadherin single-molecules in the cells under shear stress moved more in the direction of the external flow (Fig. 2, C, lower).

Next, we examined the displacement distribution of T-cadherin single-molecules in the direction of flow. The distance moved in the flow direction between each frame was measured from the tracking data of T-cadherin single-molecules in the control cells or the cells under shear stress (Fig. 2, D). Positive values for the distance represent movement in the flow direction, while negative values represent movement in the direction opposite to the flow direction. The displacement distribution of T-cadherin single-molecules was appeared as a mountain-shaped histogram centered at around zero for both the control cells and the cells under shear stress (Fig. 2, D). Compared to control cells, the histogram of the cells under shear stress showed a lower peak and broader displacement distribution, which may be due to increased membrane fluidity caused by shear stress [25, 26]. Strikingly, the average velocity for the cells under the shear stress at 20 dynes/cm^2^ was 117.5 nm/s, which is higher than the value for control cells (-27.71 nm/s). Our results suggest that the external flow drags T-cadherin anchored to the cell membrane toward the direction of flow.

### Shear stress-induced gradient distribution of transmembrane proteins

We examined whether the extracellular flow induces gradient distribution of transmembrane proteins in addition to GPI-anchored proteins. We observed XTC cells expressing the insulin receptor, Integrin ß1, E-cadherin and VE-cadherin fused with EGFP at their C-termini, which are the cytoplasmic portion of these proteins. The localization of Insulin receptor-EGFP and Integrin ß1-EGFP to the cell membrane was confirmed by single-molecule observation using Total Internal Reflection Fluorescence (TIRF) microscopy (data not shown). E-cadherin-EGFP and VE-cadherin-EGFP localized to cell-cell adhesion sites when expressed in *Xenopus* epithelial A6 cells (data not shown).

We found that applying shear stress at 10-30 dynes/cm^2^ induced the formation of a gradient distribution of Insulin receptor-EGFP and Integrin ß1-EGFP (Fig. 3, A-D, Suppl. Figs. S4 and S5). Shear stress also induced a concentration gradient distribution of E-cadherin (Fig. 3, E and F), but the ratio of the E-cadherin fluorescent intensity at the downstream region to those at the upstream region was smaller than the ratio of other membrane proteins. For Insulin receptor, the ratio of the average value within 10 µm from the downstream cell edge to the average value within 10 µm from the upstream cell edge was 0.932 for before, and, 2.64 and 2.69 for 3 and 5 min after application of shear stress at 20 dynes/cm^2^, respectively. Similarly, for Integrin ß1, the ratio of the average fluorescence intensity at the downstream region to those at the upstream region was 0.948 for before, and, 2.04 and 2.37 for 3 and 5 min after application of shear stress at 20 dynes/cm^2^. In contrast, for E-cadherin, the ratio of the average fluorescence intensity at the downstream region to those at the upstream region was 0.948 for before, and, 1.65 and 1.91 for 3 and 5 min after application of shear stress at 30 dynes/cm^2^. Taken together, the time and degree of shear stress-induced concentration gradient formation differed for each membrane protein.

**Fig. 3.**
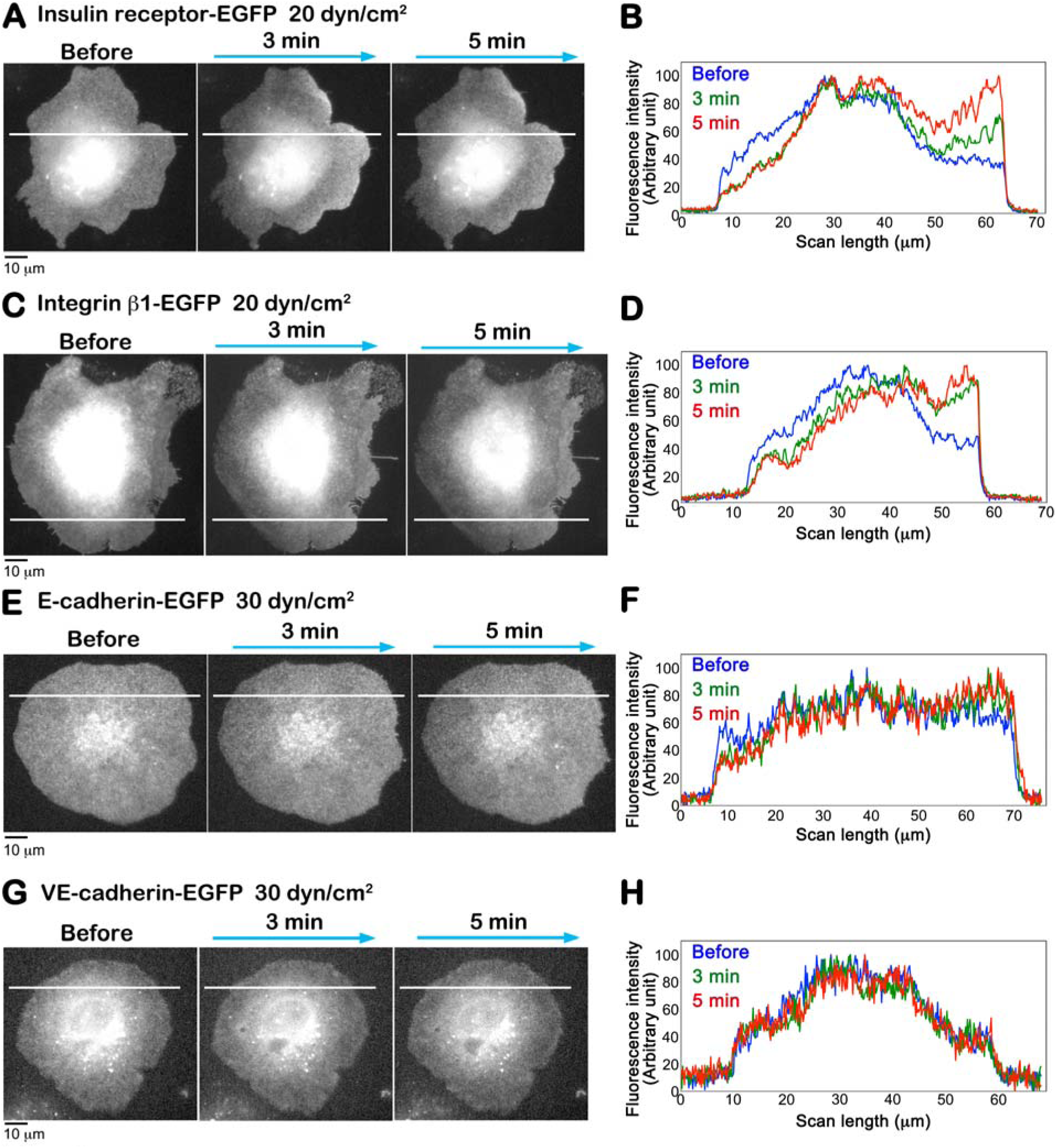
Shear stress-induced gradient distribution of transmembrane proteins. A, C, E and G, The localization of indicated proteins expressed in XTC cells, before and after the application of shear stress. B, D, F and H, Linescan analysis performed along the white lines in A,C, E and G. External flow induced gradient distributions of Insulin receptor-EGFP (A and B), Integrin β1-EGFP (C and F) and E-cadherin-EGFP (E and F), whereas VE-cadherin showed little change (G and H). A-H, The experiments were repeated at least 3 times and representative data are shown. Bars = 10 μm

In contrast to these transmembrane proteins and GPI-anchored proteins, VE-cadherin-EGFP did not form a gradient distribution 5 min after applying shear stress at 30 dynes/cm^2^ (Fig. 3, G and H). Since the cytoplasmic domain of VE-cadherin binds to the actin cytoskeleton, we examined whether disruption of the actin cytoskeleton by latrunculin B treatment enhances a gradient distribution of VE-cadherin under shear stress. However, applying the external flow did not cause any change in the distribution of VE-cadherin in the cells treated with latrunculin B (data not shown). Thus, the non-response of VE-cadherin’s distribution to shear stress cannot be explained by the linkage between VE-cadherin and the cortical actin cytoskeleton.

### Shear stress-induced gradient distribution of membrane proteins in COS-7 cells and HUVECs

We examined whether the extracellular flow induces gradient distribution of membrane proteins in mammalian cell lines, African green monkey kidney fibroblast-like COS-7 cells and human umbilical vein endothelial cells (HUVECs). COS-7 cells expressing EGFP-tagged glypican, which is a GPI-anchored protein, were exposed to shear stress. We found that applying shear stress at 17 dynes/cm^2^ induced the formation of a gradient distribution of Glypican-EGFP after 3 min and 5 min (Fig. 4, A and B).

**Fig. 4.**
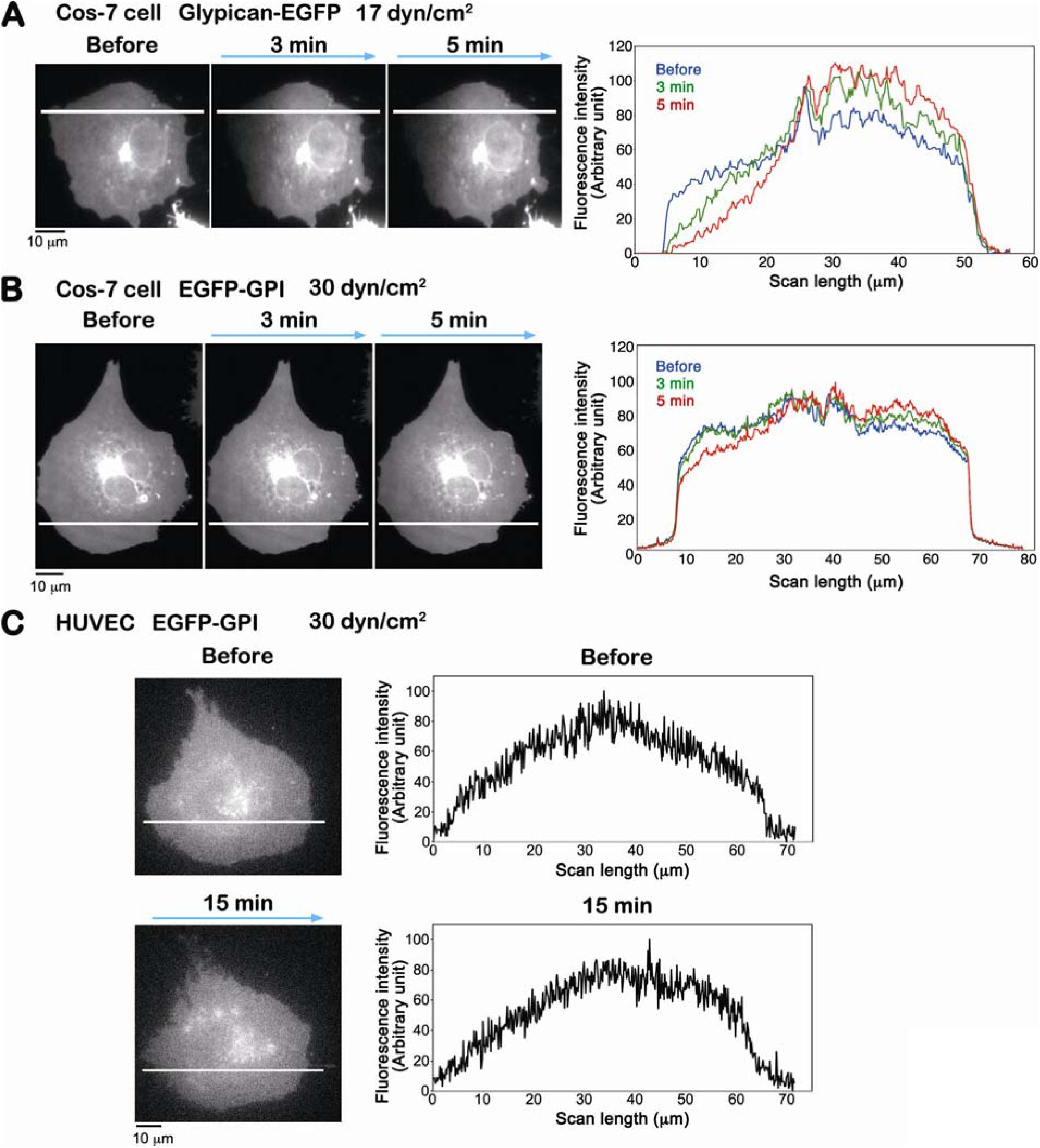
Shear stress-induced gradient distribution of GPI-anchored proteins in mammalian cells. A and B, The localization of indicated proteins expressed in COS-7 cells, before and after the application of shear stress (left images). The right graphs show the linescan analysis performed along the white lines indicated in the left images. C, The localization of of EGFP-GPI in Human umbilical vein endothelial cells (HUVEC), before and after the application of shear stress (left images). The right graphs show the linescan analysis performed along the white lines indicated in the left images. A-C, The experiments were repeated at least 3 times and representative data are shown. Bars = 10 μm

Compared to Glypican-EGFP, the shear stress induced gradient formation of EGFP-GPI occurred with a greater applied force in COS-7 cells. EGFP-GPI formed a gradient distribution 5 min after applying shear stress at 30 dynes/cm^2^ in COS-7 cells (Fig. 4, B). Similarly, in HUVECs, we observed that applying shear stress at 30 dynes/cm^2^ induced the formation of a gradient distribution of EGFP-GPI at 15 min after applying shear stress (Fig. 4, C). These results indicate that the formation of a gradient distribution of EGFP-GPI requires greater magnitude of shear stress when expressed in COS-7 cells and HUVECs than when expressed in XTC cells. Therefore, shear stress induced gradient distribution of membrane proteins in mammalian cell lines. In addition, our results suggest that the effect of shear stress on the membrane proteins varies among the cell lines.

## Discussion

Our study has revealed that both transmembrane proteins, including the insulin receptor, integrin ß1 and E-cadherin, as well as GPI-anchored membrane-linked proteins, including GPI-EGFP, T-cadherin and glypican, establish concentration gradients corresponding to the direction of external flow. By using fluorescence single-molecule analysis of gradient-forming T-cadherin, we found that T-cadherin molecules diffuse with drift in the direction of flow under shear stress. Our results suggest that the external flow directly transports a variety of membrane proteins, leading to the formation of concentration gradients.

We found that membrane proteins with structural variability establish gradient distributions under shear stress, and that the extent of gradient formation varies between membrane proteins in XTC cells (Figs 1 and 3). In the model system of supported lipid bilayers, the flow mobility of lipid-anchored proteins depends on its size and folded shape [16]. Proteins anchored to supported lipid bilayers tend to be transported more readily by flow as their molecular weight increases. Additionally, the more the proteins protrude above the bilayer surfaces in shapes that impede flow, the more efficiently they are transported by flow [16]. As shown in the flow-mediated transport of lipid-anchored proteins *in vitro*, the intrinsic characteristics of membrane protein extracellular domains likely influence the extent of shear stress-induced transport on cell membranes. We observed that shear stress causes a slight gradient formation of E-cadherin and almost no formation of VE-cadherin gradient in XTC cells (Fig. 3). The extracellular domains of E-cadherin and VE-cadherin are 80-90 kDa and rod-like structures composed of five tandem extracellular cadherin (EC) repeats. For E-cadherin, the domain has a length of approximately 18 nm and a diameter of 2 nm [27]. In solution, the E-cadherin extracellular domain monomer appears flexibly twisted under high-speed atomic force microscopy [28]. This flexible rod-like structure of E-cadherin may be less susceptible to flow forces. The extracellular domain of the insulin receptor consists of two α-subunits and parts of two ß-subunits linked by disulfide bonds, with a molecular weight of 300-400 kDa [29, 30]. Integrins are heterodimers of α and ß subunits, and the molecular weights of the extracellular domain is 210-290 kDa [31]. The extracellular domains of the insulin receptor and integrins are likely to be more susceptible to flow effects compared to E-cadherin and VE-cadherin.

The formation of shear stress-induced gradients of membrane proteins is presumably influenced not only by their intrinsic characteristics but also by their interactions with surrounding membrane components. Lateral diffusion of membrane proteins is influenced by a variety of factors, including the actin-based membrane skeleton, protein-protein interaction and protein-lipid interaction [32–34]. In particular, GPI-anchored protein motion could be slowed by extracellular proteins and proteoglycans, which might serve as obstacles or might shield the protein from flow. Also, variations in cholesterol and other lipid components across different cell types alter the viscosity of the plasma membrane, which may affect the mobility of membrane proteins. We observed flow transport of the same protein in three different cell types in order to directly compare the cortex mechanical properties. When we applied identical shear stress to GPI-anchored EGFP in XTC, COS-7, and HUVEC cells, the resulting gradient was steepest in XTC cells (Figs 1 and 4). Flow transport across the surface of COS-7 and HUVEC cells was less efficient.

The cytoplasmic domains of integrins, E-cadherin and VE-cadherin connect to actin filaments through adaptor proteins, which may restrict flow-induced transport by anchoring them to the cortical actin network [31, 35]. We tested whether disrupting the actin cytoskeleton by latrunculin B enhances VE-cadherin gradients under shear stress, but flow had no effect on the distribution of VE-cadherin in latrunculin B-treated cells (data not shown). For VE-cadherin, the linkage between its cytoplasmic domain and the actin cytoskeleton appears to make only a limited contribution to resisting external flow. The intracellular domain of the insulin receptor binds to the insulin receptor substrates but not to the cortical actin network, and therefore is unlikely to have a significant influence on extracellular flow [30].

Taking advantage of direct viewing of single molecules, we found that T-cadherin molecules exhibit diffusing motions with a bias of the flow direction. There have been few examples of the fluorescent single-molecule live-cell imaging under shear stress. In the previous study [36], external flow causes the single-nanoparticle-labeled *Clostridium perfringens* _ε_-toxin (CP_ε_T) receptor to move in the flow direction in MDCK cells, whereas when the flow is stopped, the nanoparticle-labeled receptors return to their original positions. Turkcan et al conclude that the CP_ε_T receptor is tethered to the elastic membrane structures. In contrast to the CP_ε_T receptor, T-cadherin molecules we observed in this study are presumably not tethered to the membrane structure and exhibited the diffusion mode.

VE-PTP is a receptor-type protein tyrosine phosphatase that is predominantly expressed in vascular endothelial cells [37, 38]. The previous study [19] reports that shear stress induces the redistribution in which VE-PTP accumulate in the downstream region of cultured mouse endothelial bEnd.3 cells and HEK293 cells. The redistribution of VE-PTP shown by Mantilidewi *et al* is probably similar to the flow-induced gradient formation that we demonstrate in this study. Mantilidewi *et al* conclude that shear stress-induced integrin-mediated activation of Cdc42 leads to reorganization of the actin cytoskeleton, which then causes the redistribution of VE-PTP [19]. On the other hand, we reveal that external flow directly transports membrane proteins in the flow direction, which differs from the previous VE-PTP study. Interestingly, the polarized distribution of VE-PTP accumulating downstream in the bloodstream has been demonstrated in endothelial cells in mouse thoracic aorta [20]. The downstream accumulation of VE-PTP in aortic endothelial cells suggest that shear stress-induced gradient formation of membrane proteins occurs *in vivo* as well as in cultured cells.

What is the physiological function of the shear stress-induced membrane protein gradient formation? Various type of cells, including endothelial cells, fibroblasts and neutrophils, sense external flow and exhibit responses such as morphological changes and migration along the flow direction [39–41]. Our previous *in vitro* studies demonstrated that application of low shear stress at 0.1-2 Pa (1-20 dynes/cm^2^) generates concentration gradients of lipid-anchored proteins across supported lipid bilayers [15, 16]. Based on these studies we propose that shear-stress induced membrane protein gradient formation may act in cellular sensing of flow direction [17]. Here, we demonstrate that varied membrane proteins in living cells exhibit gradient formation induced by 10-30 dynes/cm^2^ of shear stress. Application of shear stress activates various signaling pathways over seconds to minutes. For example, in endothelial cells, application of shear stress at 12 dynes/cm^2^ promotes phosphorylation of Akt, Grb2-associated binder 1 (Gab1) and endothelial Nitric Oxide synthase (eNOS) within 2 min, and the phosphorylation are peaks at 5-15 min [42]. Membrane receptors upstream of such signals may form a shear stress-induced concentration gradient, which may initiate polarized signaling in the cell. We propose that the formation of a molecular gradient along the external flow acts as a transduce, converting mechanical information of shear stress including flow direction and magnitude into polarized chemical signaling within the cell.

## Acknowledgements

This work was supported by Takeda Science Foundation (S.Y.), by Uehara Memorial Foundation (N.W.) and by Japan Society for the Promotion of Science KAKENHI Grant Number JP22H04843, JP23H04310, and JP24H01944 (S.Y.) and JP22H00456 (N.W.). Additional support came from the National Institutes of General Medical Sciences of the United States National Institutes of Health under award no. R01GM143320 (A.H.S, D.T) and from the Charles E. Kaufman Foundation (A.H.S. , D.T).

## Materials and Methods

### Plasmids

The expression vector of GPI-EGFP was obtained from Addgene (pCAG:GPI-GFP, #32601 Addgene) (Rhee et al., Genesis, 2006). The pCMV-SPORT6-*Xenopus tropicalis* T-cadherin expression plasmid was obtained from Horizon Discovery (#7658828), and the EGFP cDNA was inserted at the 5’ end of T-cadherin cDNA in the pCMV-SPORT6-T-cadherin plasmid. To generate the SNAP-tagged T-cadherin expression plasmid, the sequence encoding SNAP-tag was amplified by PCR using pSNAPf vector (New England Biolabs) and inserted into the pCMV-SPORT6-T-cadherin plasmid between the 4th and 5th T-cadherin EC domains (between amino acid 577 and 578 of *Xenopus tropicalis* T-cadherin). *Xenopus tropicalis* integrin β1 cDNA was obtained from Open Biosystems (Gene ID:394767) and was subcloned into the expression pEGFP-N3 vector harboring delCMV [43]. Mouse E-cadherin cDNA and mouse VE-cadherin cDNA were gifts from Masatoshi Takeichi (Riken, CDB, Japan), and the cDNAs were subcloned into the pEGFP-N3 delCMV vector. Two human insulin receptor cDNA fragments were fused to generate the full-length insulin receptor cDNA by using PCR. The human insulin receptor cDNA containing 5’-cDNA fragment (HsCD00021529) was obtained from the DNASU Plasmid Repository (Arizona State University). The human insulin receptor cDNA containing 3’-cDNA fragment (ORK05520) was obtained from the Kazusa DNA Research Institute (Chiba, Japan). The full-length human insulin receptor cDNA was subcloned into the pEGFP-N3 delCMV vector. A glypican-1-GFP construct with EGFP inserted into a mid-sequence disordered region was designed as described elsewhere (Sasidharan et al, personal communication) and cloned into a pcDNA3.1(+) vector by GenScript Inc (Piscataway, NJ, USA).

### Cell culture and transfection

*Xenopus laevis* XTC cells were maintained as described previously [43, 44]. Human umbilical vein endothelial cells (HUVECs) were obtained from PromoCell (Heidelberg, Germany) and cultured in an incubator at 37°C and 5% CO_2_. HUVEC cells were grown in the Endothelial Cell Growth Medium 2 (PromoCell) containing SupplementMix (PromoCell) and 1% penicillin-streptomycin (Nacalai Tesque). COS-7 cells were cultured in Cytiva HyClone Dulbecco Modified Eagle Medium (DMEM) with high glucose, L-glutamine, and sodium pyruvate, and supplemented with 10% fetal bovine serum, 100 units/mL penicillin, and 0.1 mg/mL streptomycin.

The expression plasmids were transfected into XTC cells and HUVECs by using the Neon transfection system (Thermo Fischer Scientific). COS-7 cells were transferred to DMEM lacking FBS and antibiotics. They were then transfected using Lipofectamine 2000 Transfection reagent (Invitrogen) according to the manufacturer instructions.

### Shear stress

XTC cells were seeded on a poly-L-lysine-coated μ-Slide [^0.5^ Glass Bottom slide (Ibidi) in 70% L15 Leibovitz medium (Invitrogen) without serum, riboflavin, and phenol red. Laminar shear stress was applied to the cells by running the medium through a flow channel in a μ-Slide [^0.5^ Glass Bottom slide connected by tubing a medium reservoir and a gear pump (ISMATEC). Shear stress was calculated according to the manufacturer’s instruction of the μ-Slide [^0.5^ Glass Bottom slide with the dynamical viscosity of 0.0089 dyn·s/cm^2^ for the medium without serum. The magnitude of shear stress was controlled by flow rate. HUVECs were seeded on a fibronectin-coated μ-Slide [^0.5^ Glass Bottom slide (Ibidi) in the Endothelial Cell Growth Medium 2 (PromoCell). For HUVECs, shear stress was applied in the same manner as for XTC cells at 37 °C. COS-7 cells were grown on a coated coverslip coated with poly-L-lysine which was assembled into a flow chamber (Bioptechs) just before imaging. A flow gasket in the chamber formed a microchannel with approximate height 105 µm and width 1600 µm. Shear stress was applied using a syringe pump (Harvard Apparatus) to move flow buffer (10 mM HEPES, 135 mM NaCl, 5 mM KCl, 10 mM glucose, 1 mM MgCl_2_, 1.5 mM CaCl_2_, 1 weight % fetal bovine serum and 0.5 weight % bovine serum albumin at pH 7) through the channel. The flow chamber and buffer were maintained at 37 °C throughout the experiments.

### Fluorescence microscopy and data analysis

Time-lapse imaging of XTC cells and HUVECs was acquired by using an inverted epi-fluorescent microscope (IX71, Olympus) equipped with 100-W mercury illumination, an EMCCD camera (iXon Ultra 894, Andor), and a PlanApo 1.40 numerical aperture (NA) 100× oil objective (Olympus). Images were acquired using the Metamorph software (Molecular Devise). Images were acquired at 5-s intervals for the duration of 10 to 20 mins, and shear stress was applied to cells during the time-lapse imaging. COS-7 cells were imaged using a spinning disk confocal microscope (Intelligent Imaging Innovations). Using a 63× objective, stacks were collected at a vertical spacing of 0.27 microns and excitation with a 488 nm laser at intervals of 2-4 minutes before and during flow. The stacks were converted to sum projections before linescan analysis.

Linescan analysis was performed using the Linescan function in the Metamorph software (Molecular Devices). The background intensity in an area outside of the cell was subtracted from the original image before the fluorescence intensity measurement.

### Fluorescence single-molecule imaging of SNAP-tagged T-cadherin and data analysis

XTC cells expressing SNAP-tagged T-cadherin were dissociated by trypsinization, collected by centrifugation, and then washed with serum-free 70% L15 medium. The XTC cells were collected again as a pellet and then resuspended with serum-free 70% L15 medium. SNAP-surface 549 (NEW ENGLAND BioLabs) was added to the XTC cell suspension to a final concentration of 0.2 μM and mixed gently for 15 sec. The XTC cells were collected again as a pellet and then washed with serum-free 70% L15 medium more than 3 times. The XTC cells were seeded on a poly-L-lysine-coated μ-Slide [^0.5^ Glass Bottom slide (Ibidi) in 70% L15 Leibovitz medium (Invitrogen) without serum, riboflavin, and phenol red.

Fluorescence single-molecule imaging of SNAP-tagged T-cadherin was performed using an inverted epi-fluorescent microscope (IX71, Olympus) as described in the *Fluorescence microscopy and data analysis* section. A series of fluorescence single-molecule SNAP-tagged T-cadherin images were acquired twice, before and 1 min after the application of shear stress of 20 dyn/cm^2^, for 20-40 at 100-200 ms intervals.

Single-molecule signals of SNAP-tagged T-cadherin were analyzed by using Speckle TrackerJ software as described previously [45–47]. Immobile signals were excluded from the analysis because they were stuck to the glass coverslip, and signals with diffuse behavior were tracked and analyzed.

