## supplemental information for "Shear Stress Induces Concentration Gradient Distributions of Membrane Proteins in Live Cells"

**Supplementary Materials**

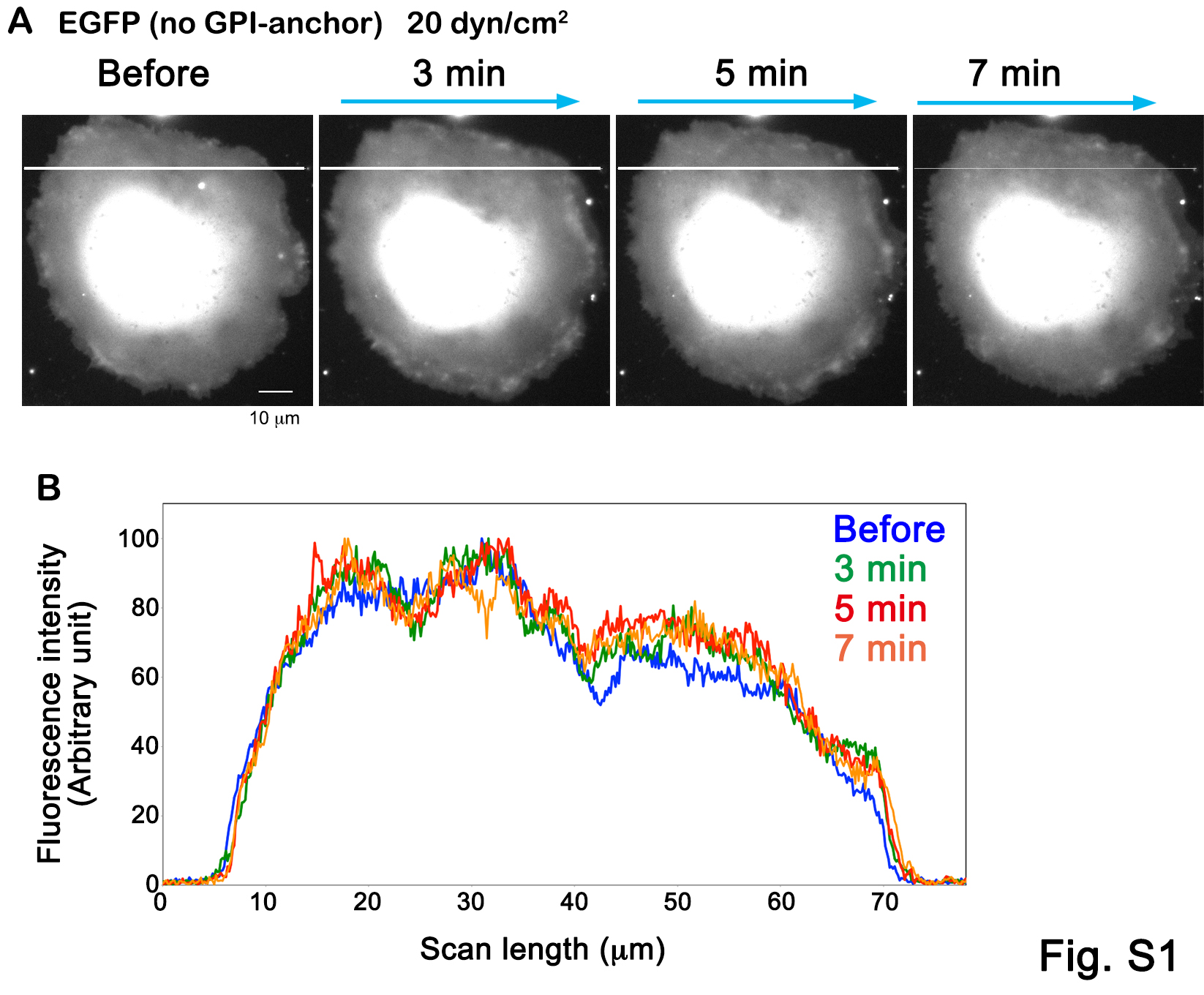

**Supplemental Fig. 1:**

Extracellular flow does not change the distribution of EGFP (no GPI-anchor). A, The localization of EGFP expressed in an XTC cell, before and after the application of shear stress in the direction indicated by arrows. B, Linescan analysis performed along the white lines in A. Bar = 10 μm

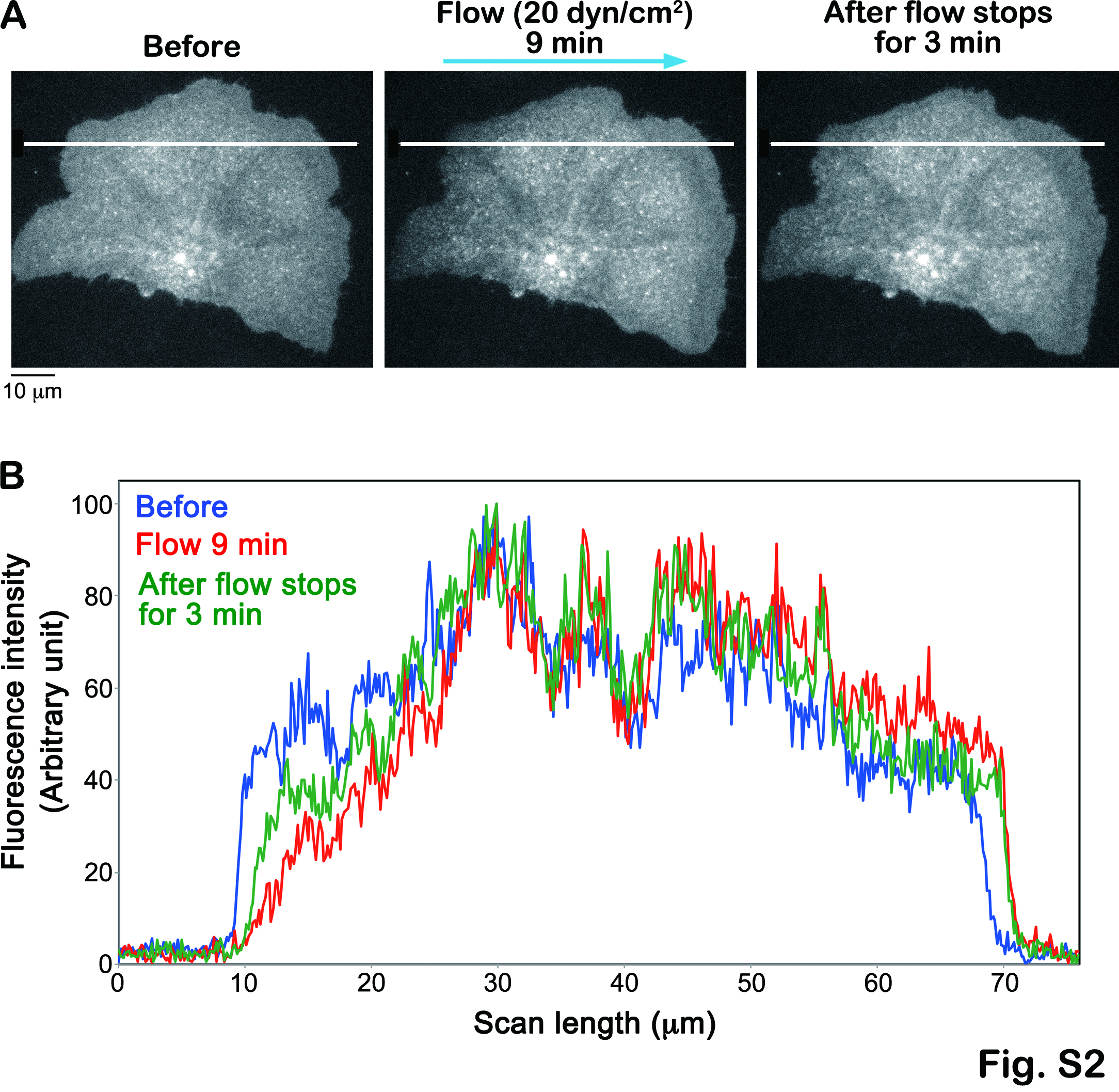

**Supplemental Fig. 2:**

After stopping the extracellular flow loading, the shear stress-induced gradient distribution of EGFP-GPI is mitigated. A, The localization of EGFP-GPI expressed in an XTC cell, before, 9 min after the application of shear stress in the direction indicated by arrows, and 3 min after stopping the flow application. B, Linescan analysis performed along the white lines in A. Bar = 10 μm

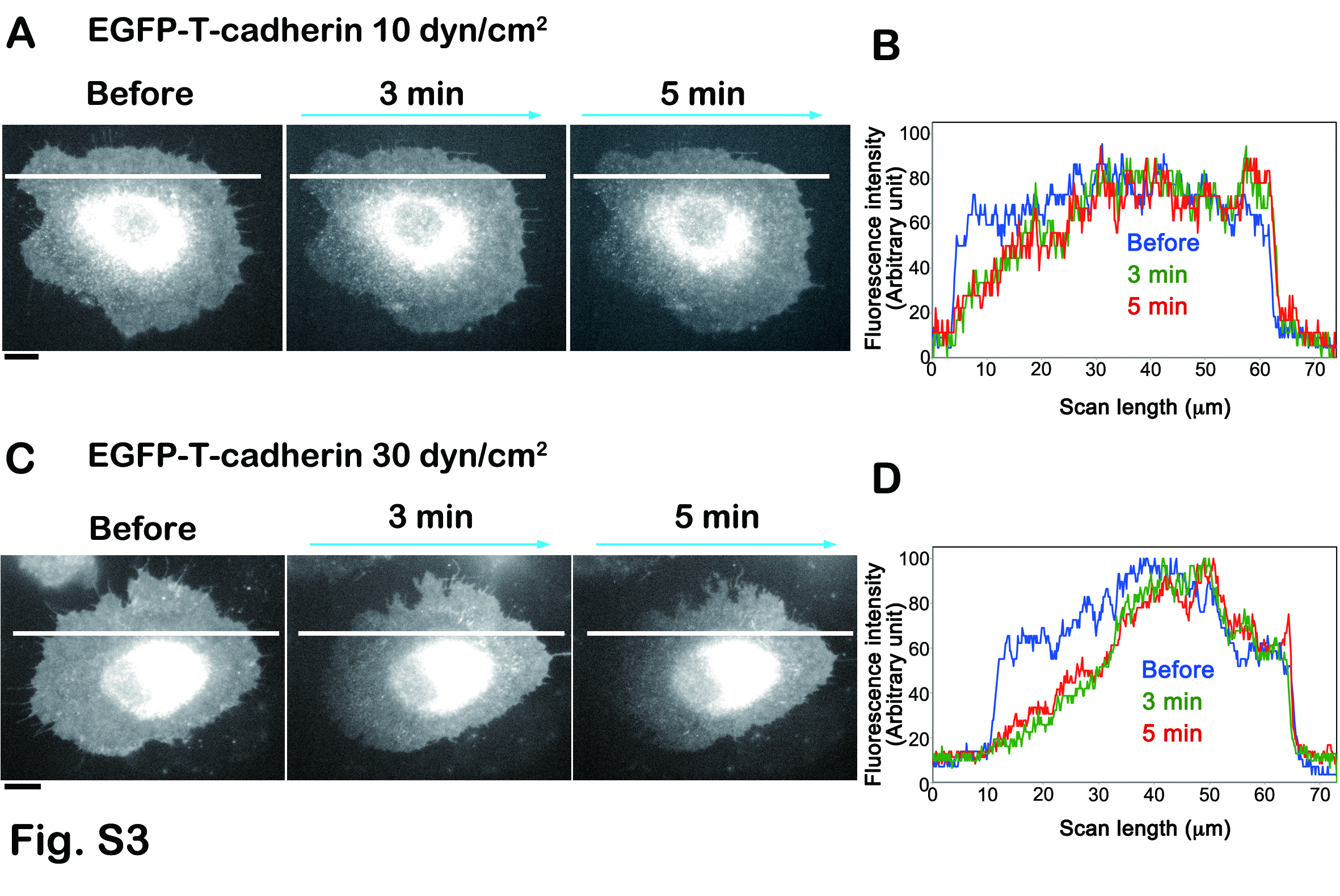

**Supplemental Fig. 3:**

The formation of EGFP-T-cadherin concentration gradient under shear stress at 10 dyn/cm^2^ (A and B) and 30 dyn/cm^2^ (C and D) in XTC cells. Bars = 10 μm

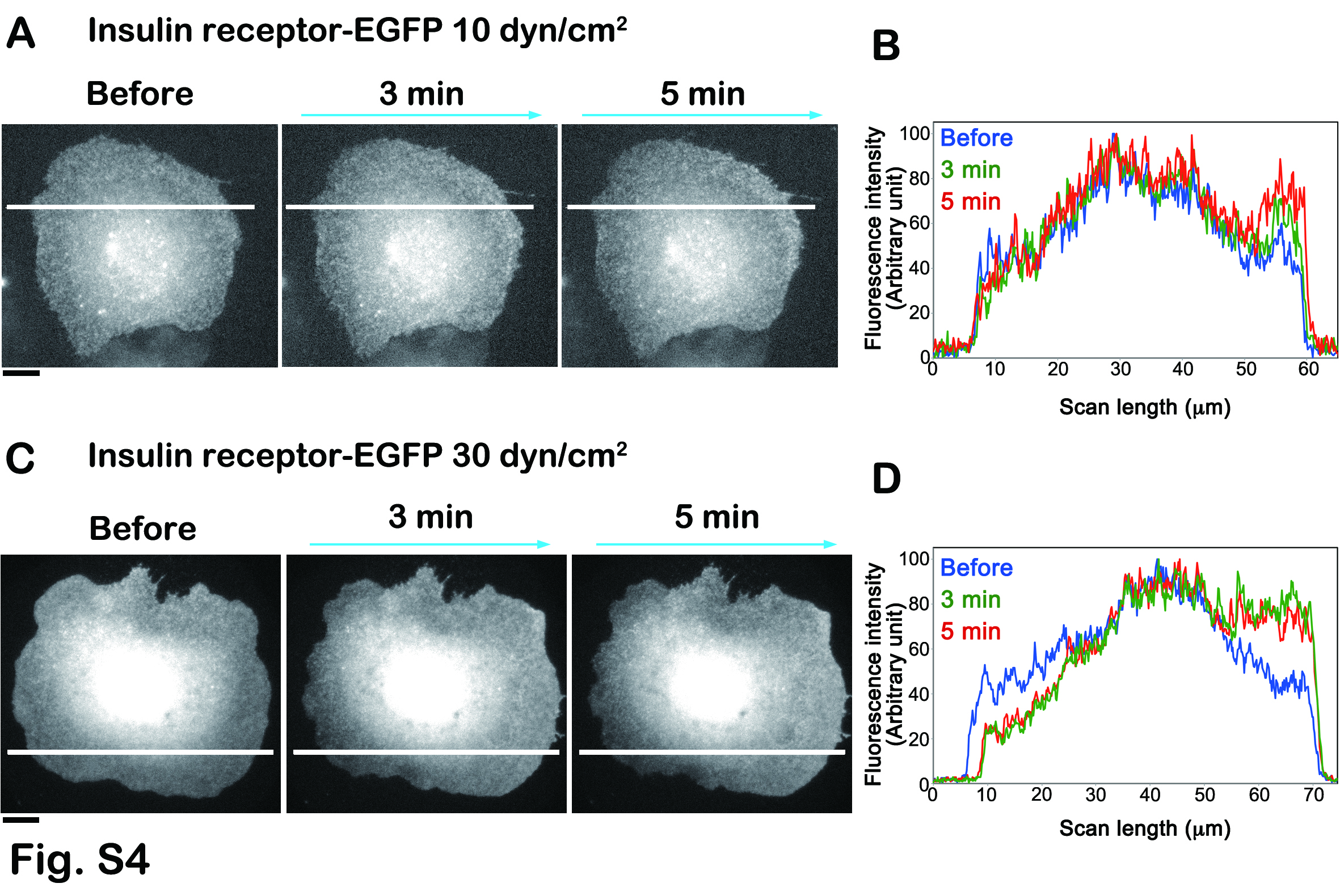

**Supplemental Fig. 4:**

The formation of Insulin receptor-EGFP concentration gradient under shear stress at 10 dyn/cm^2^ (A and B) and 30 dyn/cm^2^ (C and D) in XTC cells. Bars = 10 μm

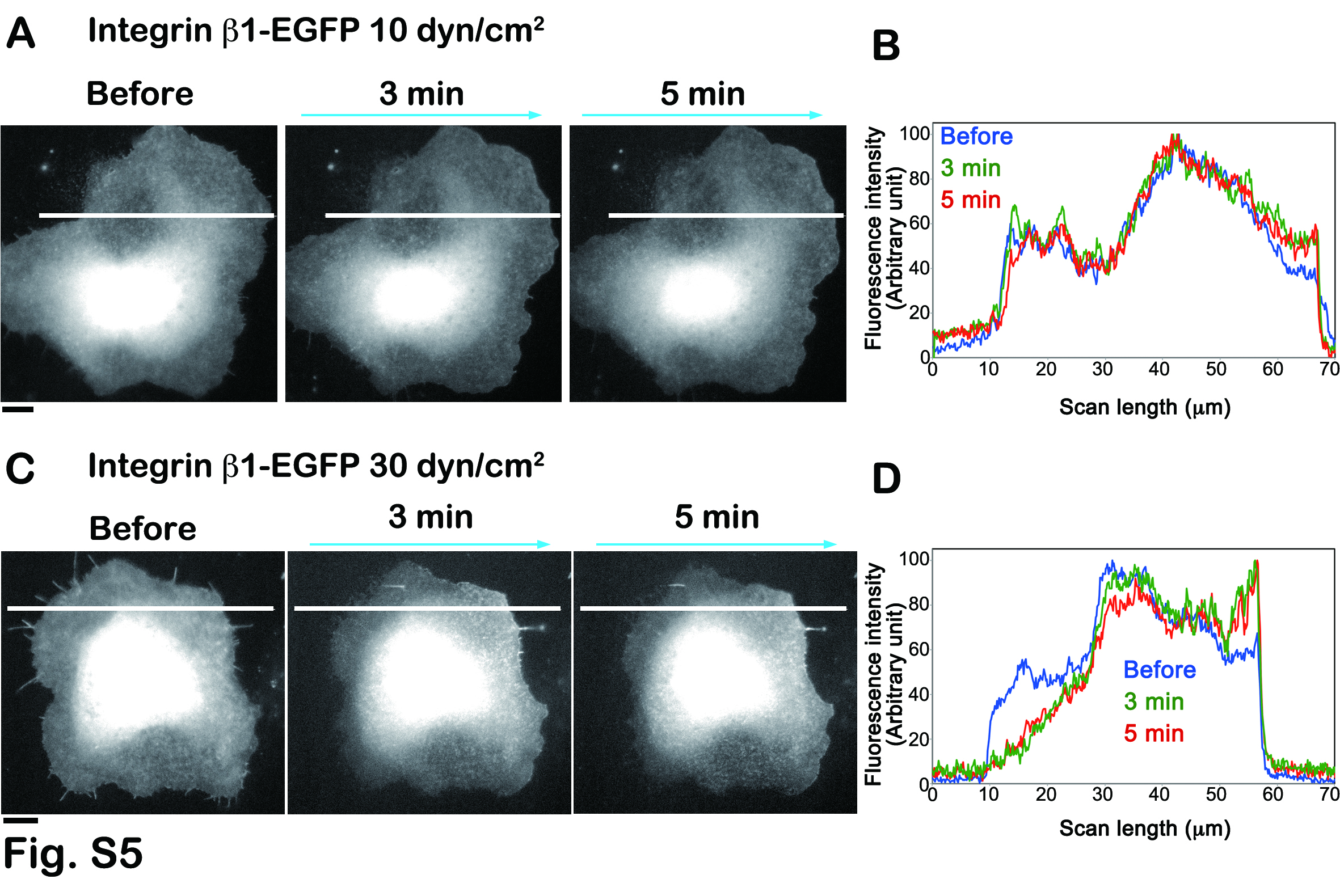

**Supplemental Fig. 5:**

The formation of Integrin β1-EGFP concentration gradient under shear stress at 10 dyn/cm^2^ (A and B) and 30 dyn/cm^2^ (C and D) in XTC cells. Bars = 10 μm

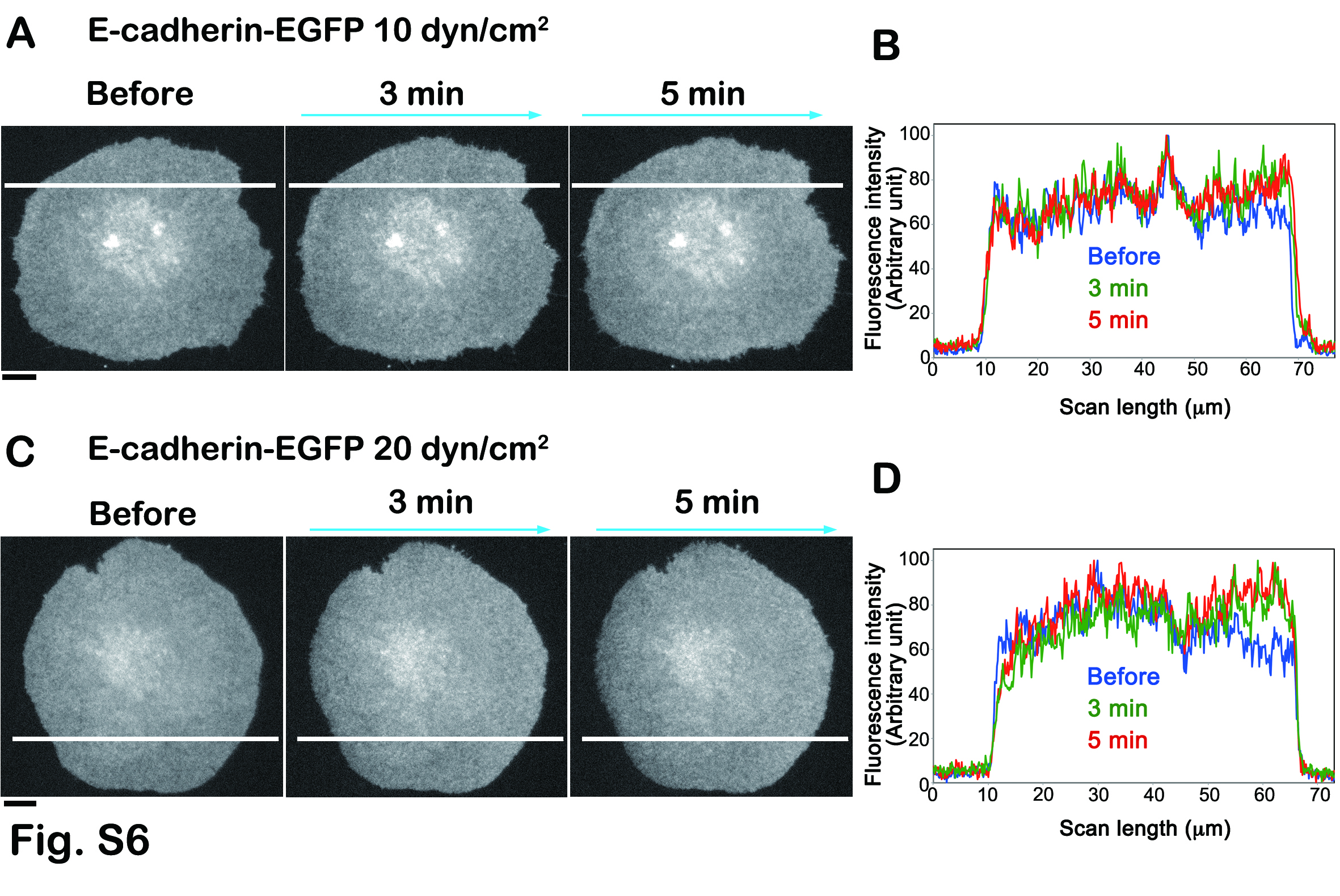

**Supplemental Fig. 6:**

The formation of E-cadherin-EGFP concentration gradient under shear stress at 10 dyn/cm^2^ (A and B) and 20 dyn/cm^2^ (C and D) in XTC cells. Bars = 10 μm
